# Yeast MoClo secretion and surface display toolkit 2.0: improvements and applications for analysis of protein-protein interactions and whole-cell biocatalysis

**DOI:** 10.1101/2025.08.19.671047

**Authors:** Vanja Jurić, Leah G. Erwin, Nicola M. O’Riordan, Eamonn Maher, Justin D. Holmes, Paul W. Young

## Abstract

*Saccharomyces cerevisiae* is an invaluable model organism for both fundamental biological research and biotechnological applications including recombinant protein production as well as protein and metabolic engineering. We previously developed a modular cloning (MoClo) based toolkit for *S. cerevisiae* that facilitates rapid optimization of signal peptides and anchor proteins for efficient secretion and/or surface display of heterologous proteins of interest. Here we describe further improvements and applications of this yeast secretion and display (YSD) toolkit. New parts encoding anchor proteins based on N-terminal fusion to a truncated Aga1 and C-terminal fusion to Aga2, each with three possible epitope tag options, are described. We also added parts that facilitate high throughput detection of secreted proteins of interest through GFP fluorescence complementation and parts encoding “secretion boosting” yeast proteins, whose overexpression has previously been reported to enhance secretion of heterologous proteins. In addition, two surface display applications of the toolkit are showcased. We demonstrate that yeast surface display of an anti-GFP nanobody allows cost-effective evaluation of the interactions of GFP-tagged proteins of interest, either by flow cytometry or yeast-based co-immunoprecipitation. In addition, using yeast cells as whole-cell catalysts, we show that co-display of the poly(ethylene terephthalate) (PET) degrading enzyme leaf-branch compost cutinase with hydrophobin1 enhances the breakdown of PET plastic, while triple co-display of these proteins with MHETase causes complete conversion of the intermediary monohydroxyLethyl-terephthalate (MHET) to terephthalic acid. The diverse applications described herein demonstrate the broad applications of the updated MoClo YSD toolkit 2.0 in both synthetic biology and other research fields.

## INTRODUCTION

The invention and continuous improvement of methods for DNA synthesis and assembly underpins a lot of synthetic biology research, in many cases facilitating the “build” step of the Design–Build–Test–Learn experimental cycle that is much vaunted in the field. Beyond these methods themselves, broader hierarchical frameworks are required to simplify, standardize and accelerate the development of complex, multicomponent synthetic biological systems. Modular cloning (MoClo) is a successful example of such a framework that employs type IIS restriction enzymes to enable efficient assembly of plasmids containing single or multiple transcriptional units (*1*). This general MoClo strategy has been widely adopted (*2, 3*). The MoClo yeast toolkit (MoClo YTK) developed by Dueber and coworkers provided highly characterised parts such as promoters, terminators and selectable markers (*4*). This toolkit has gained traction among synthetic biologists working with *S. cerevisiae* and other yeast species and has spawned several extensions – compatible toolkits that share the same “syntax”. Examples of this are expansions with greater functionality for CRIPSR-based applications (*5–7*), genomic integration (*7, 8*), intracellular protein targeting (*9*) and G-protein coupled receptor engineering (*10*) in *S. cerevisiae,* as well as toolkits for the yeasts *Komagataella phaffi* (*11*)*, Kluyveromyces marxianus* (*12*)*, Schizosaccharomyces pombe* (*13*) and even insect and mammalian cells (*14, 15*). Such toolkits provide the means to rapidly and reliably design and build large numbers of single or multigene expression constructs that can then be evaluated and iteratively optimized to achieve the desired performance of the synthetic biological system that is being engineered.

Yeast surface display is a highly versatile technique that is widely used in both biotechnology and fundamental research contexts. It is based on the simple concept of retaining a protein of interest (POI) on the outer surface of yeast cells by expressing it fused to a yeast cell wall anchor protein. The most common yeast display system employs the Aga1p and Aga2p cell wall proteins (*16*). Aga1p has a glycosylphosphatidylinositol (GPI) anchor that tethers it to the yeast cell membrane, while Aga2p binds Aga1p through two disulfide bonds. A heterologous POI can be fused to either the amino or carboxyl (N- or C-) terminus of the Aga2p protein (*17*). Subsequent to the development of this system, other *S. cerevisiae* cell wall proteins, such as Cwp2p, Tip1p Flo1p, Pir1p-4p and Sed1p, have been adapted as alternative surface display anchors and synthetic anchor proteins have also been developed (*18–20*). Inclusion of an epitope tag on the anchor protein allows the efficiency of surface display of a POI to be assessed quantitatively by flow cytometry. Yeast display is most widely used as a screening platform to either identify proteins with a desired property or to optimize specific characteristic of a lead protein. Proteins libraries with ∼ 10^8^ variants can readily be screened using magnetic or fluorescence-activated cell sorting (FACS) or other selection strategies. The properties of interest might be binding parameters in the case of antibody or non-antibody related affinity reagents, the activity or substrate specificity of enzymes, or protein stability (*17*).

A distinct application of yeast display is the generation of whole-cell biocatalysts by immobilizing one or more enzymes on the yeast cell surface (*18, 20, 21*). This approach avoids the need to purify enzymes and provides for easy recovery and reuse of enzymes for multiple rounds of catalysis. Other potential advantages compared to soluble enzyme systems may include enhanced enzyme stability, control of the protein orientation, co-display of several enzymes to mimic supramolecular complexes and the possibility of shuttling reaction products generated extracellularly into intracellular metabolic pathways. One example of such yeast display-based biocatalysis is bioethanol production from cellulosic biomass by co-displaying cellulolytic, amylolytic and xylanolytic enzymes (*18, 22*). Another is the generation of a glucose biosensor using yeast displaying glucose oxidase (*23*).

Regardless of the specific experimental goal, careful consideration of numerous factors is required to achieve optimal yeast cell surface display of a POI. These include the choice of promoter, signal peptide, display anchor, host strain and culture conditions (*24*). Systems that standardize and streamline the selection and optimization of these parameters have great potential to accelerate yeast display-based research. The MoClo framework is an ideal platform for rapid optimization of expression constructs and host strains for yeast surface display. However, a comprehensive plasmid toolkit for this purpose has not been available to date.

We previously developed the yeast secretion and display modular cloning (MoClo YSD) toolkit for *S. cerevisiae* that facilitates rapid optimization of signal peptides for efficient secretion and/or surface display of heterologous POIs (*25*). Five anchor proteins were included in this toolkit and their basic function when used in conjunction with a panel of signal peptide containing parts was characterized. Here we describe further improvements to this toolkit, especially the yeast display aspect. Additional anchor proteins, including those that provide the option of C-terminal fusion of proteins of interest have been added and characterised. For most anchor proteins, parts providing a choice of three epitope tags have been included. This provides flexibility for co-display experiments as demonstrated through the generation of a whole cell biocatalyst for PET degradation that features co-display of three proteins. Novel yeast display applications of the toolkit to examine protein: protein interactions by co-immunoprecipitation and flow cytometry are also described. Parts to enable high throughput detection of secreted proteins through GFP fluorescence complementation and a panel of cDNAs that can be screened for enhancement of protein secretion or display have also been added to the toolkit. The resulting MoClo YSD 2.0 toolkit, used in conjunction with the MoClo YTK and its many compatible derivative kits, constitutes a comprehensive resource to design, build and optimize yeast-based systems that involve secretion and/or surface display of proteins.

## RESULTS AND DISCUSSION

### Expansion of the MoClo YSD toolkit to facilitate co-display and carboxyl-terminal fusion of displayed proteins

We previously generated MoClo parts encoding five different HA epitope-tagged yeast display anchors that facilitate fusion of displayed protein at the N-terminus of the anchor protein (*25*). Of these, Aga1/Aga2, Sed1, and the synthetic anchor 649 stalk performed best. Parts encoding different anchor proteins with different epitope tags would facilitate co-display of multiple proteins. We therefore generated 2xFLAG and MYC tagged versions of the Sed1 and 649 stalk anchors, which are utilised later in this study to display PETase enzymes. One limitation of the classical Aga1/Aga2 system is the need to co-express both proteins – ideally at a 1:1 molar ratio. To avoid this complication we generated MoClo compatible type 4a parts encoding HA, 2xFLAG and MYC epitope-tagged versions of a truncated Aga1 protein (Aga1ΔN181) that was described previously by Yang and coworkers(*26*). Used within the framework of the YSD toolkit, these parts allow proteins of interest and signal peptides, encoded as type 3b’ and 3a’ parts respectively, to be expressed as amino-terminal (N-terminal) fusions to Aga1ΔN181. To test the Aga1ΔN181 anchors we displayed an anti-GFP Nanobody (α-GFP Nb) (*27*) on yeast cells and evaluated binding of GFP as well as anti-tag antibodies by flow cytometry (Figure 1A). For the HA and 2xFLAG tagged constructs, greater than 80% of yeast cells were labelled with both GFP and anti-tag antibodies. For the MYC-tagged anchor GFP labelling was again > 80%, but labelling with an anti-MYC antibody was slightly lower at ∼70% and exhibited more variability compared to other anti-tag antibodies.

**Figure 1.**
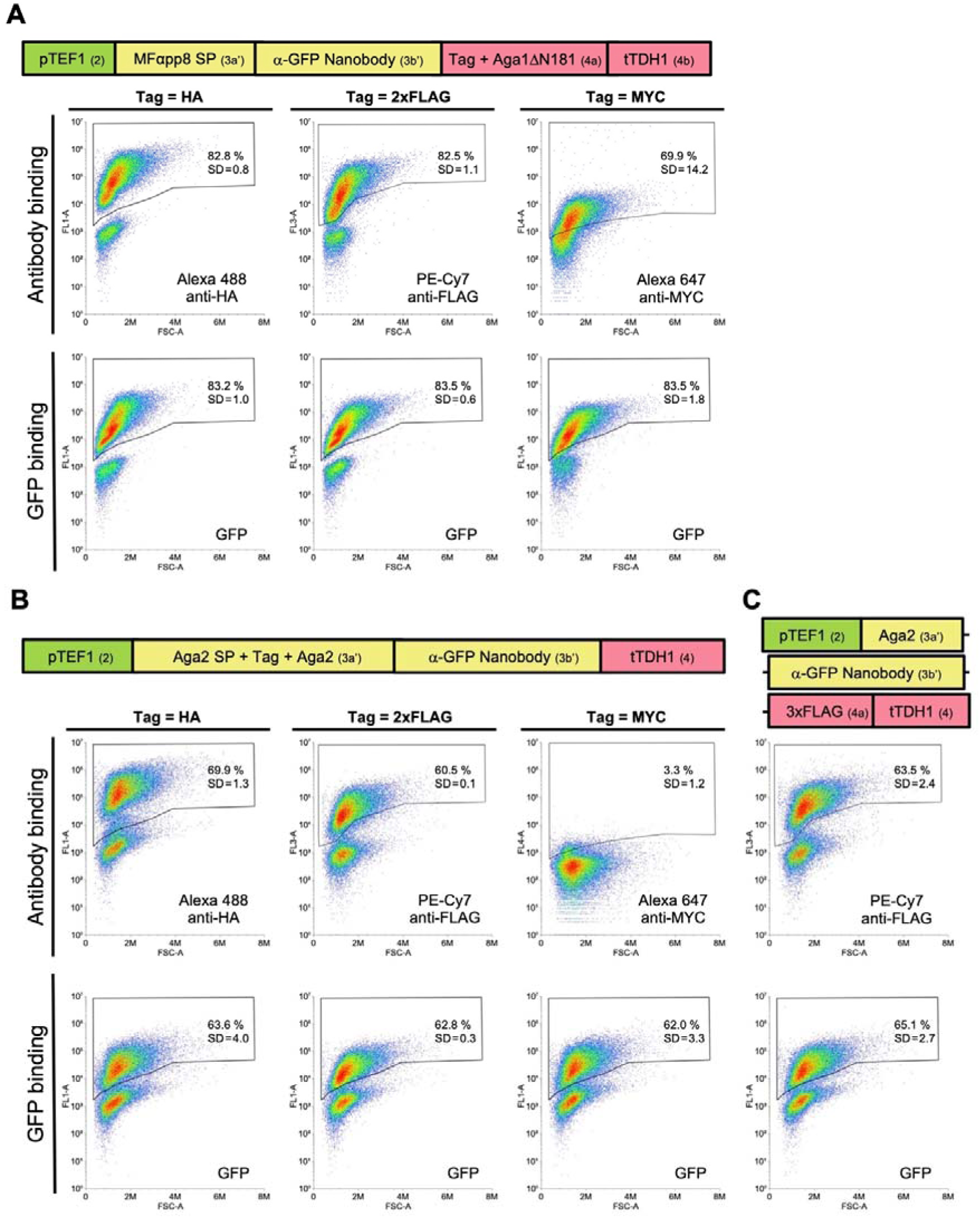
Characterization of MoClo compatible Aga1 and Aga2 based yeast surface display anchors with HA, FLAG and MYC epitope tags. **(A**) Type 4a MoClo parts encoding an N-terminally deleted Aga1 protein lacking the first 181 amino (ΔN181) with three different epitope tags were assembled into expression constructs that display the α-GFP Nb as an N-terminal fusion. Surface display was assessed by anti-tag antibody binding, while the functionality of the nanobody was indicated by binding of GFP from a bacterial cell lysate. **(B,C)** Type 3a’ MoClo parts encoding Aga2 with each of three epitope tags after the signal peptide (B) or an untagged Aga2 (C) were assembled into expression constructs that display the α-GFP Nb as a carboxyl terminal fusion when co-transformed with a plasmid expressing full length Aga1. Surface display and functionality of the nanobody was assessed as in (A). Plots depict the relevant fluorescence signal (Y-axis) versus forward light scattering (X-axis). The mean percentage of positively stained cells from three independent experiments is shown on a representative plot with the standard deviation (SD) indicated. Schematics of the MoClo-generated expression constructs indicate the promoter, terminator and signal peptide (SP) sequences used as well as part types (in brackets).

**Figure 2.**
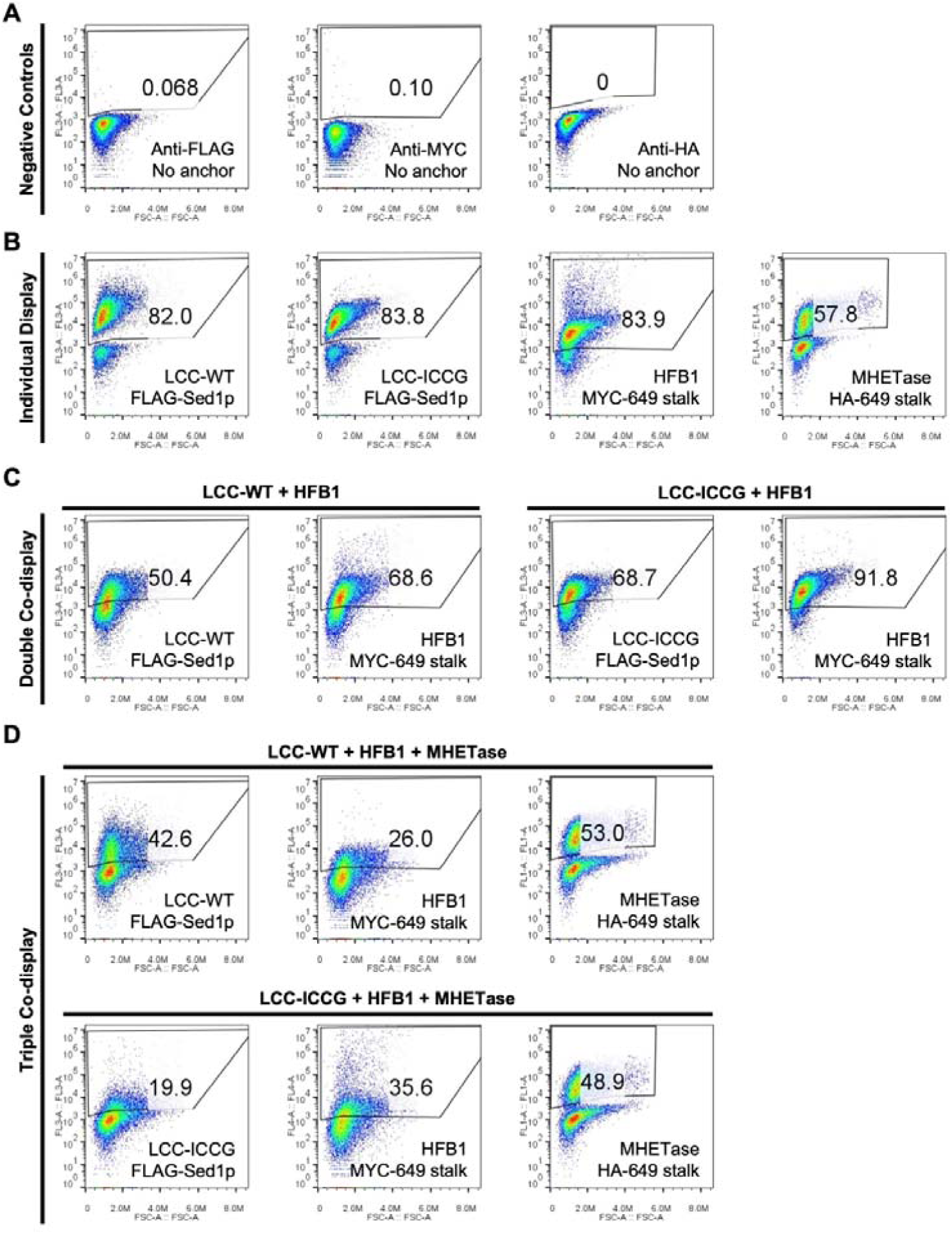
Yeast surface display and co-display of LCC-WT, LCC-ICCG, HFB1 and MHETase using different combinations of display anchors and SPs. **(A)** The indicated proteins were expressed in yeast cells in combination with the indicated display anchors. Anchors with either a 2xFLAG-tag, MYC-tag or HA-tag were used and surface display was assessed by flow cytometry using PE-Cyanine7 anti-FLAG, Alexa-647 anti-MYC and Alexa-488 anti-HA antibodies respectively. Plots depict the relevant fluorescence signal (Y-axis) versus forward light scattering (X-axis), with the percentage of positively stained cells indicated. A representative example from 2-3 independent experiments is shown in each case. **(A)** Untransformed yeast cells stained with each antibody were used as negative controls. **(B)** Proteins displayed individually using the OST1-pre SP for LCC-WT and LCC-ICCG, and the MFαpp8-pre-pro SP for HFB1 and MHETase. **(C)** Co-display of LCC-WT or LCC-ICCG with HFB1 using the same SPs as in (B). **(D)** Triple co-display of the LCC variants, HFB1 and MHETase with the Aga2 SP used for MHETase in this case. Further details of expression plasmids can be found in Table S1.

**Figure 3.**
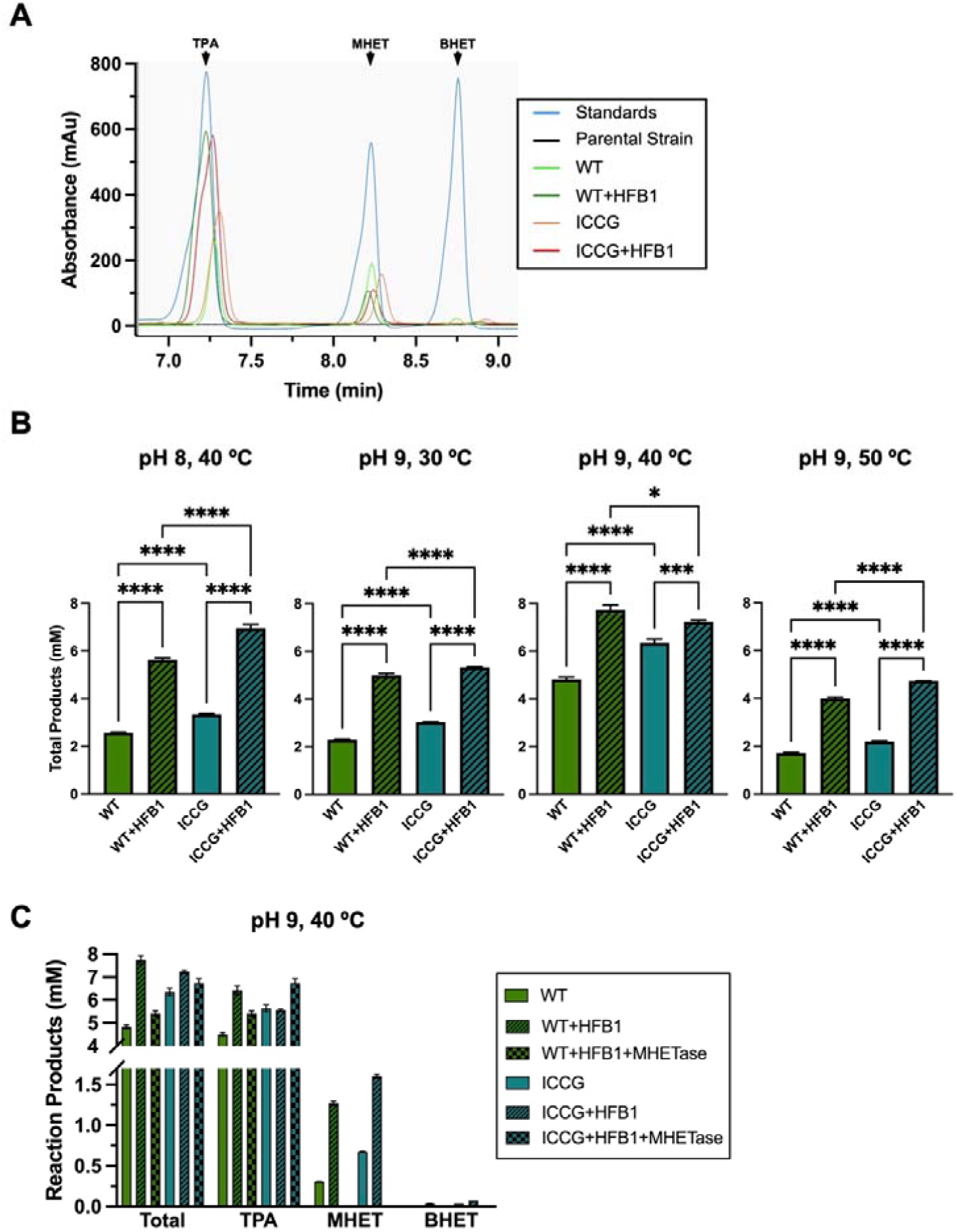
Quantitative analysis of the breakdown of low-crystallinity PET by yeast strains displaying combinations of LCC enzymes, HFB1 and MHETase. **(A)** Representative chromatograms from HPLC analysis of PET degradation products produced in 24 hours by yeast strains displaying the wild type (WT) or ICCG variant of LCC +/-HFB1 at 50 °C and pH 9. Chromatograms for the TPA, MHET and BHET standards used to generate the standard curve have also been overlaid on the graph. The area under the curve was used to determine the concentration of each product. **(B)** Quantification by HPLC of total products (TPA + MHET + BHET) released over 3 days by strains displaying either the WT or ICCG variant of LCC +/-HFB1 at the indicated combinations of temperature and pH. **(C)** Quantification by HPLC of total products and a breakdown of individual products released over 3 days at pH 9, 40 °C by the strains shown in (A) and (B), in addition to those displaying MHETase in combination with LCC and HFB1. Mean values from three samples tested per condition are shown with error bars representing the standard deviation.

The ability to fuse a protein of interest to the C-terminus of a display anchor is desirable in situations where N-terminal fusion interferes with the function or expression of the displayed protein. This is not possible with most anchor proteins as their C-terminus is attached to the yeast plasma membrane via a GPI lipid anchor. However, fusion to the C-terminus of Aga2 has been widely used in the classical Aga1/Aga2 system (*17*). To add this capability to the YSD toolkit we generated MoClo compatible type 3a’ parts encoding untagged, as well as HA, 2xFLAG and MYC epitope-tagged versions of the Aga2 protein. These parts lack a stop codon and have a linker sequence that permits fusion to a POI encoded by a type 3b’ part.

Aga2 expression constructs with the α-GFP Nb as POI were co-expressed with full-length Aga1 and tested by flow cytometry. Greater than 60% of yeast cells expressing the HA, 2xFLAG tagged and untagged Aga2 proteins were positively labelled by both GFP and anti-tag antibodies (Figure 1B, 1C). By contrast, while GFP binding to the α-GFP Nb expressed as a fusion with MYC-tagged Aga2 was very efficient (> 60% positive cells), detection using an anti-MYC tag antibody was extremely poor (∼3% positive cells). As a demonstration of the utility of this system we employed it to observe a PDZ domain: ligand interaction. PDZ domains typically bind a C-terminal sequence on their interaction targets making C-terminal fusion of this binding motif essential to maintain functionality. Specific binding of YFP-tagged Erbin PDZ domain to an Erbin PDZ binding motif (*28*) displayed as a C-terminal fusion with 2xFLAG Aga2 could readily be detected by flow cytometry (Figure S1A).

Using the Aga1ΔN181 anchor, the α-GFP Nb is displayed with similar efficiency to what we previously observed using Aga1/Aga2, Sed1 or the synthetic 649 stalk anchors (*25*). This adds another highly efficient anchor with a choice of three epitope tags to the YSD toolkit.

The Aga1/Aga2 display system requires co-expression the two proteins – ideally at a 1:1 molar ratio, since excess Aga2 will be secreted from the cell in the absence of binding via disulphide bonding to Aga1. The Aga1ΔN181 anchor circumvents this consideration and also uses up just one selectable marker, which may be an important consideration for co-display experiments depending on the auxotrophy of the host strain being utilized.

The flexibility to fuse a protein of interest via either N or C-terminal fusion to an anchor protein can be highly desirable in yeast surface display experiments, particularly when trying to preserve native enzymatic or binding activity (*24*). Here we have added this advantageous feature of the Aga1/Aga2 system to the YSD toolkit. While untagged Aga2 and all three epitope tagged versions of Aga2 seem to effectively display the α-GFP Nb fused to their C-termini, detection of the MYC tag with an antibody was very inefficient. This suggests that the MYC epitope is either inaccessible or perhaps undergoes proteolysis in the context in which we have added it between the signal peptide the sequence of the mature Aga2 polypeptide. Pir proteins have also been used as surface display anchors that facilitate C-terminal fusions (*29*). We developed MoClo-based Pir2 anchors and used them to display the α-GFP Nb (Fig S1B). However, their surface display could not be detected via epitope tags and they showed very modest binding to GFP. It is possible however that they would work better with a different POI and thus have been added to the toolkit. While we primarily used the α-GFP Nb to characterize these new anchors in this study, both the Aga1ΔN181 and C-terminal Aga2 anchors show efficient display of the fluorescent protein mRuby2 (*30*) (Figure S1C) – a protein whose secretion is very dependent on the SP used (*25*). With the addition of these new parts, the YSD toolkit, when used in conjunction with the original MoClo YTK, represents the most comprehensive and flexible system for the design and optimization of yeast surface display constructs that we are aware of.

### Co-display of leaf-branch compost cutinase with hydrophobin 1 and MHETase to generate a whole-cell biocatalyst for plastic degradation

There is significant interest in enzymatic approaches for recycling or valorising plastic waste. In particular, numerous enzymes that can break down PET plastic have been identified and in some cases engineered to enhance their catalytic properties (*31–34*). One potential strategy for scaling up such approaches is through the development of whole-cell biocatalysts that avoid the need to extract and purify enzymes (*20*). Cell surface display of PETase enzymes provides access to solid plastic substrates and offers the potential for separation and reuse in multiple rounds of catalysis (*35*). One of the most promising PETase enzymes identified to date is leaf-branch compost cutinase (LCC) (*36*), that has been improved through directed evolution to produce variants such as LCC-ICCG that has increased hydrolysis activity and thermostability (*37*). To our knowledge, yeast surface display of LCC enzymes has not been described to date. We therefore sought to apply the YSD toolkit for surface display of LCC and the LCC-ICCG variant while also co-displaying hydrophobin 1 (HFB1). HFB1 is a small fungal protein from *Trichoderma reesei* that mediates attachment to hydrophobic surfaces and has previously been shown to enhance substrate attachment and PET degradation activity of *Ideonella sakaiensis* PETase when co-displayed on the surface of the methylotrophic yeast *Komagataella phaffii (P. pastoris)* (*38*). In addition, we wanted to examine the feasibility of triple co-display of LCC enzymes and HFB1 with *I. sakaiensis* MHETase which converts the intermediary PET breakdown product monohydroxyLethyl terephthalate (MHET) into terephthalic acid (TPA) and ethylene glycol (*39*).

We first displayed each protein individually using different combinations of signal peptides and anchor proteins from the YSD toolkit. Efficient display of each protein was achieved as assessed by flow cytometry. Greater than 80% of cells displaying LCC, LCC-ICCG and HFB1 were labelled with an antibody recognizing the epitope tag on the anchor protein, while for MHETase this figure was 57.8%. When the LCC variants were co-displayed with HFB1 the percentage of cells labelled positively for each protein generally decreased but was still greater than 50% in all cases. Triple co-display of the LCC variants with HFB1 and MHETase resulted in a further decrease in display efficiency, but between ∼20 and 53% of cells were still positive labelled for each protein.

Next we examined the ability of these yeast strains to degrade low-crystallinity PET discs using HPLC to detect and quantify the reaction products. Firstly, wild type LCC and the engineered ICCG variant, either alone or co-displayed with HFB1 were assessed at four combinations of temperature and pH. Total products (the sum of the molar concentrations of TPA, MHET and BHET) were assessed as a measure of PET degradation. While the whole-cell biocatalysts were active at all temperature/pH combinations, the highest activity for each strain was observed at 40 °C and pH 9. The strains co-displaying HFB1 with either LCC or LCC-ICCG exhibited significantly higher activity than strains displaying the LCC enzyme alone. The magnitude of enhancement of activity varied with the reaction conditions but in many cases co-display of HFB1 with LCC doubled PET degradation compared to the strain displaying the LCC enzyme alone.

To examine the effects of co-displaying MHETase we focused on the optimal reaction conditions (40 °C and pH 9) and examined the individual reaction products. TPA was the major reaction product, followed by MHET, with only trace amounts of BHET for strains expressing either LCC enzyme, alone or in combination with HFB1. However, triple co-display of MHETase along with LCC and HFB1 resulted in complete conversion of MHET to TPA, which was the sole reaction product detectable. Total reaction products generated by the triple co-displaying strains was lower than those co-displaying LCC and HFB1 but higher than the strain expressing the LCC enzymes alone.

The MoClo YSD toolkit facilitated the rapid identification of anchor proteins and signal peptides for efficient surface display of LCC, HFB1 and MHETase. Furthermore, the results demonstrate that all three proteins are functionally active when displayed or co-displayed.

This cannot be taken for granted as another PET degrading enzyme, FAST-PETase (*40*), for which we achieved efficient surface display as assessed by flow cytometry, did not show any PET degrading catalytic activity (V.J. unpublished observations). We show that co-displaying HFB1 with the LCC enzymes on *S. cerevisiae* enhanced PET degradation, extending the previous finding of a similar effect of HFB1 on the catalytic efficiency of *I. sakaiensis* PETase when co-displayed on *K. phaffii* (*38*). This lends support to the co-display HFB1, or other proteins that enhance interactions with the surface of plastics, as a general strategy for whole-cell biocatalysts in this field.

The flexibility of the YSD toolkit allowed us to readily combine surface display of LCC and HFB1 with a third protein –MHETase. While the surface display of MHETase in *S. cerevisiae* had previously been described (*41*), combining this with co-display of LCC facilitated the conversion of PET completely into TPA and ethylene glycol, eliminating the intermediates MHET and BHET in the final product. This should greatly simplify the recycling or valorisation of PET degradation products in such a biocatalytic system. In addition, high MHET levels may cause inhibition of the PET-degrading enzymes, reducing overall catalytic efficiency (*42*). Indeed PETase-MHETase fusion proteins have been developed to address this challenge (*42*). In our system LCC and LCC-ICCG produce a relatively low proportion of MHET as reaction product compared to *I. sakaiensis* PETase (*38*) or its derivative FAST-PETase (*42*). It is not clear whether this is an intrinsic difference between these enzymes or due to differences in substrate properties such as crystallinity. In any case, MHETase co-display may be even more beneficial for these PETase enzymes or for more challenging high-crystallinity PET substrates.

Co-display or two and especially three proteins led to significant decreases in the display efficiency of a given protein compared to when it is displayed on its own. This probably explains the decrease in total products generated by the triple versus double co-displaying yeast strains. This issue likely reflects intrinsic capacity limitations of the yeast protein expression and secretion processes. The MoClo framework provides tools to potentially circumvent this problem (*1, 4*). For example, displayed MHETase hydrolyses MHET very efficiently and could potentially be displayed at significantly lower levels using a weaker promoter or less efficient display anchor without affecting TPA yield. Likewise, inducible promoters or constitutive promoters of varying strength could be used to titrate the expression and display of HFB1 and LCC (or other PETase enzymes) to achieve optimal levels. Triple co-displaying strains catalysed more PET breakdown than those solely displaying LCC despite the triple co-display strains have substantially less efficient display of the LCC enzymes as assessed by flow cytometry. This suggests that the gains in PET degradation by simply maximizing the expression and display levels of LCC may be marginal and highlight the impact of HFB1 co-display. Overall, this example demonstrates the utility of the YSD toolkit and broader MoClo framework in generating sophisticated surface display-based whole-cell biocatalysis systems.

### Utilization of yeast displaying an **α**-GFP Nb to analyse protein:protein interactions

Given the efficient GFP binding observed for yeast displaying an α-GFP Nb (α-GFP-Nb yeast) we decided to examine whether these yeast could be used to immunoprecipitate GFP or GFP-tagged proteins from cell lysates with a view to potentially examining protein:protein interactions by co-immunoprecipitation. Immunoprecipitation of GFP from a lysate of GFP-expressing *E. coli* cells by α-GFP-Nb yeast was compared to control cells displaying mRuby2. Proteins were eluted using either low pH or by boiling in protein gel loading buffer containing 4% SDS. Specific immunoprecipitation of GFP was observed by western blotting for both elution methods although the release of significant quantities of yeast proteins upon boiling in 4% SDS was apparent by Coomassie staining (Figure 4A). We tested three elution conditions similar to ones previously described (*43*) (0.2 M Glycine pH 2.5, 0.2 % SDS/100 mM Tris pH 7.5 and 8 M Urea) at five different temperatures and found that incubation of yeast cells in 8 M Urea for 5 minutes at 75 °C provided the most efficient elution of fluorescent proteins as assessed by western blot (Figure S2A). This elution condition significantly reduced the release of yeast proteins compared to boiling in 4% SDS (Figure S2B) although low pH elution at temperatures between 4 and 55 °C provided even cleaner elution of GFP with reasonable efficiency. We were curious to compare the GFP binding capacity of α-GFP-Nb yeast to the commercially available and widely used GFP-Trap reagents that are based on anti-GFP nanobodies conjugated to magnetic particles or agarose beads. Following immunoprecipitation from *E. coli* lysates, GFP was eluted either by incubation at low pH (37 °C) or in 8 M Urea (75 °C) and analysed by Western blotting (Figure 4B). Using a range of volumes of GFP-Trap elutes to generate a standard curve, the binding capacity of 1 × 10^8^ α-GFP-Nb yeast was calculated in terms of an equivalent volume of GFP-Trap beads. By this analysis the binding capacity of 1 × 10^8^ α-GFP-Nb yeast was equivalent to ∼12 ul and ∼1.2 ul of GFP-Trap magnetic agarose beads for low pH and 8 M Urea elution respectively. The difference in these values can be largely attributed to comparatively less efficient elution of GFP at low pH from the GFP-Trap beads.

**Figure 4.**
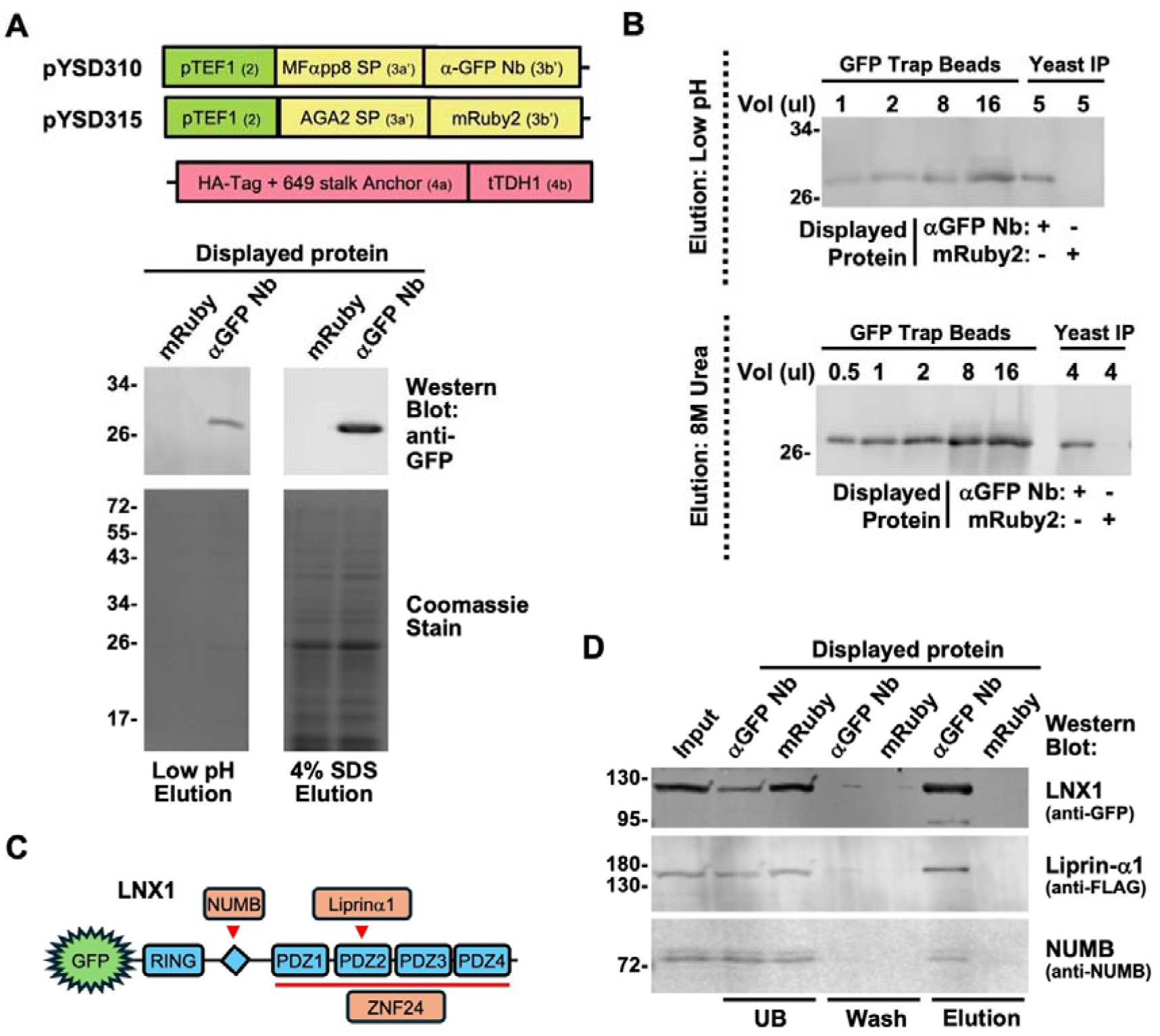
Application of the yeast displaying an α-GFP Nb to analyse protein:protein interactions by co-immunoprecipitation. **(A)** Specific immunoprecipitation of GFP using yeast. 1 × 10^8^ yeast cells displaying either an α-GFP Nb (α-GFP-Nb yeast) or mRuby2 (as a negative control) were incubated with a lysate from GFP-expressing *E. coli* cells. After washing, bound proteins were eluted either by incubating at 37 °C in 0.2 M Glycine pH 2.5 (low pH) or boiling in SDS-PAGE sample buffer (4% SDS) for 5 minutes. Eluates were analysed by western blotting and Coomassie staining to detect GFP and total protein respectively. A schematic of the MoClo-generated expression constructs used is shown indicating the promoter, terminator, SP and displayed protein as well as part types (in brackets). **(B)** Comparison of GFP binding capacity of α-GFP-Nb yeast to commercially available GFP binding (GFP Trap) magnetic agarose beads. Proteins were eluted from 7 ul GFP Trap beads or 1 × 10^8^ yeast cells by incubating for 5 minutes at low pH (as in (A)) or in 8 M Urea at 75 °C. The indicated volumes of eluate were analysed by western blotting to detect GFP. Yeast cells displaying mRuby2 served as a negative control. **(C)** Schematic representation of GFP tagged LNX1 indicating its domain structure and its previously described interacting proteins NUMB, Liprin-α1 and ZNF24. **(D)** Specific co-immunoprecipitation of GFP-LNX1 with FLAG tagged Liprin-α1 and endogenous NUMB from HEK293 cell lysates using α-GFP-Nb yeast. mRuby2 displaying cells served as a negative control. The unbound (UB), wash and low pH elution fractions are shown, with the sizes of the molecular weight markers indicated in kDa on the left.

We then examined whether α-GFP-Nb yeast could be used to study protein: protein interactions by co-immunoprecipitation. The well-characterised interactions of the E3 ubiquitin ligase LNX1 with NUMB and Liprin-α1 (*44–46*) were chosen to address this question (Figure 4C). GFP-tagged LNX1 and FLAG tagged Liprin-α1 were co-transfected into HEK293 cells and cell lysates subjected to immunoprecipitation using either α-GFP-Nb yeast or control yeast displaying mRuby2. FLAG-tagged Liprin-α1 as well as endogenous NUMB were specifically co-immunoprecipitated with GFP-LNX1 by α-GFP-Nb yeast but not control yeast cells (Figure 4D).

Next we asked whether these interactions could be observed directly by flow cytometry – avoiding the need for time consuming gel electrophoresis and western blotting. α-GFP-Nb yeast incubated with lysates from HEK293 cells transfected with either FLAG-tagged Liprin-α1 or NUMB plus either GFP or GFP-LNX1 were analysed by flow cytometry. GFP-LNX1 bound very efficiently to α-GFP-Nb yeast (>70% of cells labelled). Specific binding of FLAG-tagged Liprin-α1 and Numb to GFP-LNX1 in comparison to GFP alone was observed, with 30% and 10% of cells positively labelled with an anti-FLAG antibody respectively (Figure 5A and 5B). We also examined a less well characterized interaction of LNX1 with ZNF24 in this manner (*46*). The specific interaction of LNX1 with FLAG-ZNF24 was observed with ∼77% FLAG positive cells detected in the presence of GFP-LNX1 versus ∼3% for cells incubated with lysate containing only FLAG-ZNF24 (Figure 5C).

**Figure 5.**
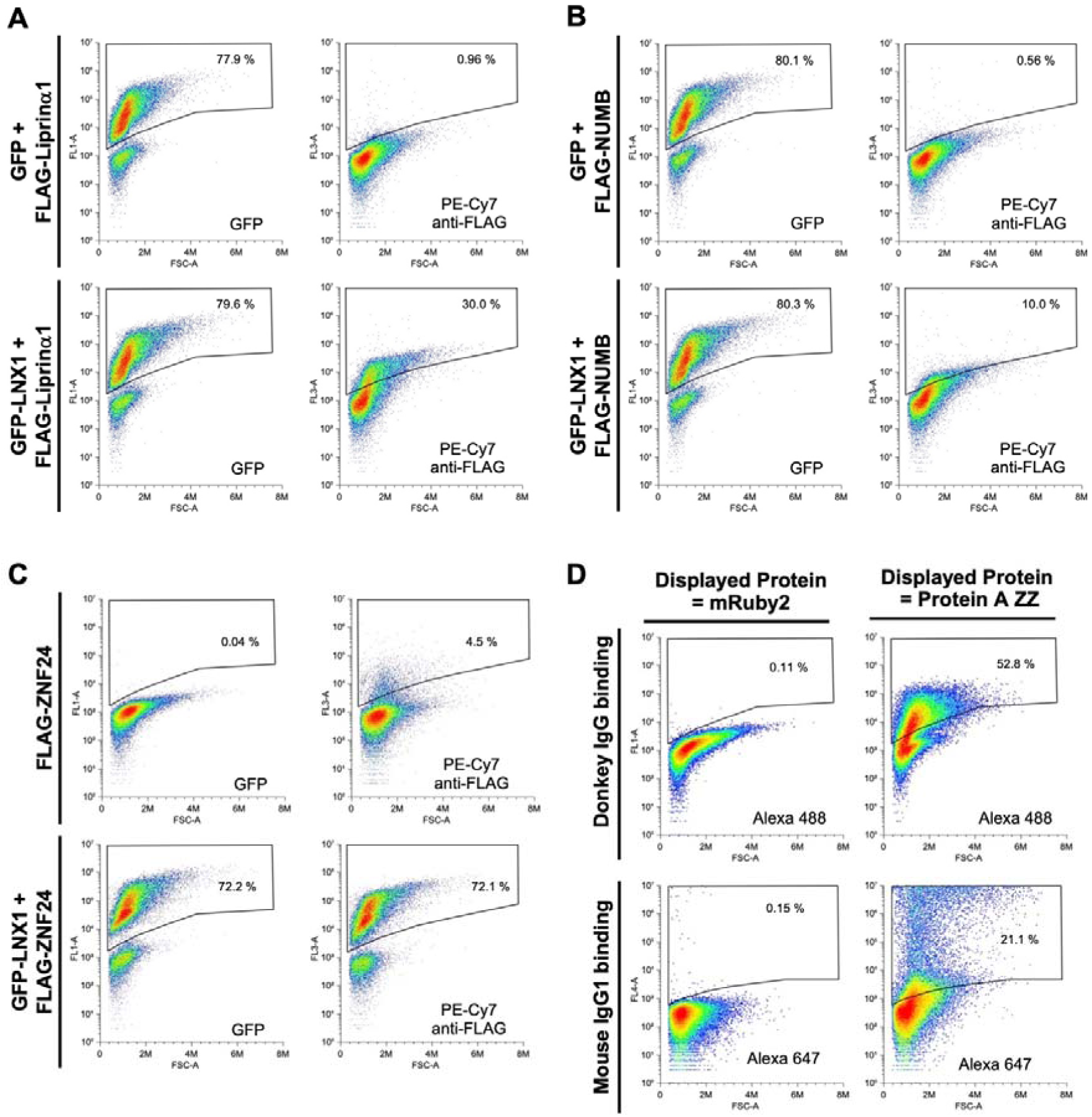
Application of the yeast displaying an α-GFP Nb or Protein A ZZ domain to analyse protein:protein interactions by flow cytometry. **(A-C)** Detection of the Liprin-α1, NUMB and ZNF24 interactions with LNX1 by flow cytometry. α-GFP-Nb yeast were incubated with lysates from HEK cells expressing the proteins indicated on the left and analysed by flow cytometry using an anti-FLAG epitope tag antibody. FLAG tagged Liprin-α1, NUMB and ZNF24 are detected bound to α-GFP-Nb yeast specifically in the presence of GFP-LNX1. **(D)** Efficient and specific capture of immunoglobulins by yeast displaying the Protein A ZZ domain. Yeast were transformed with the pYSD325 that directs surface display of the ZZ domain using the MFαpp8 SP and an HA-tagged 649-stalk stalk anchor protein. Fluorophore conjugated donkey IgG and mouse IgG1 binding to yeast displaying either mRuby2 (as a negative control) or the ZZ domain was assessed by flow cytometry. Plots depict the relevant fluorescence signal (Y-axis) versus forward light scattering (X-axis), with the percentage of positively stained cells indicated. A representative example from 2-3 independent experiments is shown in each case.

Finally, recognising that it is often preferable to study the interactions of untagged endogenous proteins that have not been over expressed, we asked whether yeast cells could be functionalized to capture antibodies that might recognise such endogenous POIs. Tandem Z domains (ZZ) based on the B domain of *Staphylococcus aureus* protein A have previously been used for this purpose (*47*). We therefore created a MoClo part encoding the ZZ domain and screened expression constructs containing several combinations of SPs and display anchor in order to develop yeast strains that gave efficient surface display (Figure S3A).

Some SPs used gave negligible surface display of the ZZ domain while others were very efficient – demonstrating the value of screening multiple SPs that is afforded by the YSD toolkit. One of the strains with good surface display could efficiently capture IgG antibodies from both donkey and mouse (Figure 5D). This opens up the possibility of employing such yeast strains to analyse endogenous protein complexes using antibodies directed against native proteins by either co-immunoprecipitation or flow cytometry based assays.

These results highlight α-GFP-Nb yeast as a potentially flexible and cost-effective reagent to study the interactions of GFP-tagged POIs. For a lab with basic microbial culturing capability, α-GFP-Nb yeast can be generated on demand with the primary cost being selective yeast growth media and labour. We estimate that a 0.6 L culture (OD_600_ = 4) will contain α-GFP-Nb yeast with a GFP-binding capacity equivalent to a 500 ul vial of commercial GFP-binding magnetic agarose beads. Considering just the price of the culture media, α-GFP-Nb yeast could be generated for less than 5% of the cost of the commercial beads. α-GFP-Nb yeast could therefore represent a very economical alternative to GFP-affinity beads in situations where cost is critical due to funding limitations or the need to analyse large numbers of samples. The possibility of examining protein: protein interactions on the surface of yeast cells by flow cytometry would be particularly advantageous for such low-cost, high-throughput analyses.

GFP is widely used to visualize proteins, especially in living cells. While it can also be used as an affinity reagent through the use of GFP binding antibodies and nanobodies, there are situations in which a smaller affinity tag is desirable. As a step towards functionalizing yeast for these purposes, we displayed an engineered anti-HA epitope tag scFv antibody (*48*) using the YSD toolkit and demonstrated that is was functional (Figure S3B). Yeast displaying the ZZ domain offer the possibility of studying protein:protein interactions using antibodies against any protein of interest – very similar to traditional co-immunoprecipitation using Protein A agarose beads, for example. However, the IgG binding profile of the ZZ domain could limit such approaches. We report efficient capture of donkey IgG and mouse IgG1 and would expect good binding to rabbit IgG (*49*). However, we saw poor binding to rat IgG2 and goat IgG (data not shown), in line with previous reports (*49*) and the known binding specificity of full-length Protein A. The display of other naturally occurring or engineered domains from Protein A, or other immunoglobulin binding proteins, would be required to circumvent these limitations. As shown for the ZZ domain (Figure S3A), the YSD toolkit is well-suited to optimize SP and anchor combinations for efficient surface display of such proteins.

Yeast display has been used previously for immunoprecipitation, primarily in order to identify and characterize antigens for displayed scFv antibody fragments (*43, 50, 51*). We are not aware of previous studies employing yeast display for co-immunoprecipitation of interacting proteins from cell lysates. The use of yeast cells for this purpose is thus novel, but of course has potential drawbacks. Interactions of POIs with the yeast cell wall could cause non-specific binding or potentially occlude binding sites on these proteins. The release of yeast proteins under more efficient, but harsher, elution conditions is potentially problematic – particularly if one wanted to study interactions of yeast proteins. However, for studying proteins for non-yeast species for which species-specific antibodies are available, or if both interacting proteins are epitope tagged, the presence of yeast proteins in eluates should not affect western blot analysis. The possibility of protein degradation by yeast-derived proteases during the immunoprecipitation procedure should also be considered, though we have not observed this problem. Growing yeast cells for each experiment is an inconvenience, however we observed that α-GFP-Nb yeast cultures stored at 4 °C retained their GFP binding capacity for at least two weeks (Figure S2C). Lyophilization may be an option for longer term storage based on studies of other surface displayed proteins (*52*).

Another consideration is that larger volume of yeast cells compared to GFP affinity beads may require more extensive washing or larger elution volumes. Nevertheless, while not suitable for every application, our findings suggest that α-GFP-Nb yeast represent a viable low-cost alternative for GFP-based co-immunoprecipitation experiments in certain contexts.

### Validation of GFP complementation as a readout of protein secretion for the YSD toolkit

The YSD toolkit provides a platform to rapidly screen sixteen signal peptides to optimize secretion (or surface display) of a protein of interest in yeast. Potentially, this might be combined with optimization of other parameters (culture conditions, promoters, coding sequence variants etc.) necessitating high-throughput analysis of protein secretion efficiency. To facilitate such multiparametric analyses we developed new parts for the YSD toolkit that allow protein secretion to be assessed by GFP fluorescence complementation in a microtiter plate format – a more high-throughput approach compared alternatives such as gel electrophoresis and western blotting. We employed the self-complementing GFP fragments GFP1-10 and GFP11-M3 (GFP11) (*53, 54*). Type 4a MoClo parts encoding the 16 amino acid GFP11 sequence, as well as GFP11 with a C-terminal polyhistidine (His^6^) tag were generated. As an initial validation of the approach we generated a yeast strain that expresses invertase with a C-terminal GFP11-His^6^ fusion (Figure 6A). Secreted invertase was captured from culture supernatants via the His^6^ tag and incubated with GFP1-10 purified from *E. coli*. The development of GFP complementation fluorescence was detectable after 90 minutes and continued to increase over 24 hours (Figure 6B). Next, to demonstrate that the system could be used to differentiate signal peptide efficiency, strains that express mRuby2-GFP11-His^6^ with three different signal peptides were evaluated. A strain expressing mRuby2-FLAG-His^6^was used as a negative control. Levels of secreted mRuby2 were determined both by measurement of mRuby fluorescence and by western blotting (Figure 6C). The Aga2 and ECM14 SPs directed mRuby2 secretion much more efficiently than CRH1, in line with our previous observations (*25*). These differences were very well correlated with measurements of GFP complementation fluorescence (Figure 6D). The control FLAG-His^6^ construct exhibited negligible fluorescence complementation despite being secreted at high levels – demonstrating the specificity of fluorescence complementation for the presence of GFP11.

**Figure 6.**
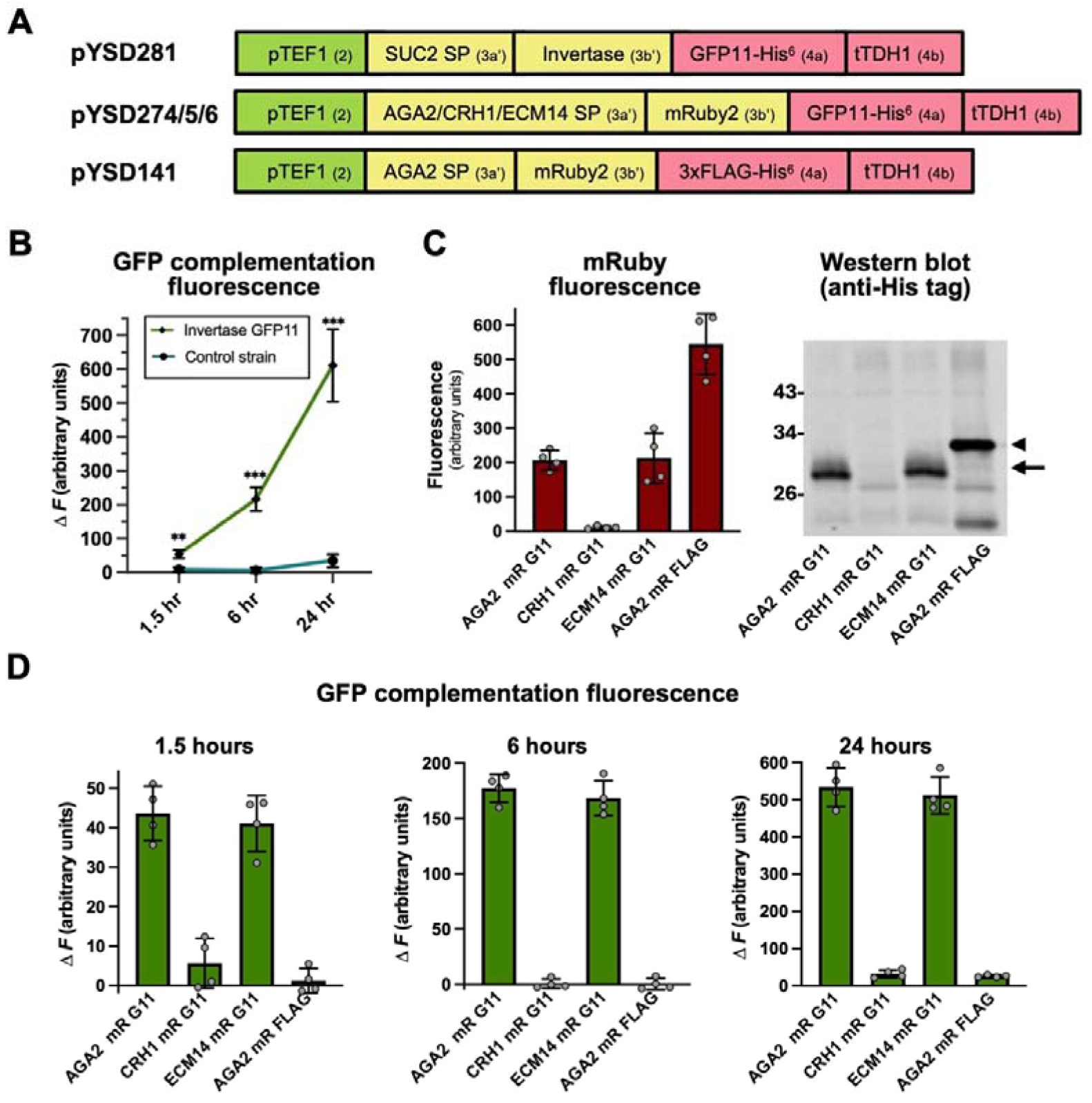
Development of GFP complementation as a readout of protein secretion for use with the yeast surface display toolkit. **(A)** Schematic representations of the expression constructs used indicating SPs, secreted POIs, and GFP11 or control detection tags as well as promoter and terminator sequences. Part types are indicated in brackets. **(B)** Secreted invertase with a carboxyl-terminal GFP11-His^6^ tag (from a yeast strain transformed with pYSD281 plasmid) was concentrated using Ni agarose resin and incubated with bacterially expressed and purified GFP1-10. GFP complementation fluorescence was measured after 1.5, 6 and 24 hours and compared to a control strain that did not secrete any heterologous protein. **(C,D)** Yeast strains that direct secretion of GFP11-His^6^ tagged mRuby2 (mR G11) utilizing three different signal peptides, as well as a control strain that secretes FLAG-His^6^ tagged mRuby2 (mR FLAG) were generated by transformation of plasmids pYSD274/5/6 and pYSD141 respectively. His-tagged mRuby2 proteins were concentrated from the media using Ni agarose resin and quantified based on mRuby2 fluorescence and western blotting (C) as well as GFP complementation fluorescence (D). The arrowhead and arrow in (C) indicate the position on the western blot of FLAG-His^6^ and GFP11-His^6^ tagged mRuby2 respectively. Graphs plot the mean fluorescence measurements from four biological replicates with individual data points and error bars representing SD shown.

With modular cloning it is easy to design and build large numbers of expression constructs for a protein (or proteins) of interest. High throughput methods are needed to test such designs in order to avoid an “evaluation bottleneck” (*55*). Fluorescence complementation has been used in other contexts for rapid evaluation of protein expression and solubility (*54, 56, 57*). Adapting it for the YSD toolkit allows protein secretion levels in yeast to be readily interrogated through fusion of the short 16 amino acid GFP11 sequence to a POI. This small tag is less likely to interfere with protein function or trafficking, or to add a substantial metabolic burden to the host cell in comparison to fusion with an intact fluorescent protein, for example. High throughput colorimetric assays can often be developed to detect a protein of interest, particularly in the case of enzymes. However, this must be done on a case by case basis and may not always be possible – necessitating detection using low throughput methods such as gel electrophoresis and western blotting. Fluorescence complementation allows the same scalable detection paradigm to be used independently of the protein being studied. The GFP11 parts developed and characterised here should therefore be a valuable addition to the YSD toolkit.

### Curation of a panel of MoClo compatible yeast coding sequences previously shown to enhance secretion of heterologous proteins

Despite significant advances in understanding and modelling the secretory pathway in *S. cerevisiae* optimising the secretion of a specific protein of interest remains challenging (*58*). A strain developed for production of one heterologous protein will not necessarily efficiently secrete a different protein. Traditional cloning methods limit the number of genetic perturbations that it is practical to screen for their ability to enhance secretion. Modular cloning approaches can potentially increase the ease with which variants of expression constructs for a protein of interest can be combined with either overexpression of a “secretion boosting” gene or deletion of a “secretion limiting” gene – facilitating rapid prototyping of yeast production strains.

With this goal in mind, we have developed a panel of twenty-four “secretion boosting” *S. cerevisiae* proteins that have been shown in previous studies to enhance the secretion of one or more heterologous POIs (Table 1). These proteins function in the different steps of the secretory pathway that may be a bottleneck that limits secretion of a specific protein. Coding sequences for these proteins were domesticated by removal of BsaI and BsmBI restriction endonuclease recognition sites (making them compatible with the yeast MoClo toolkit) and cloned into the level 1 YTK entry vector pYTK001. Single gene or multigene overexpression constructs for these “secretion booster” proteins can be rapidly assembled utilizing the wide range of transcriptional control elements and selectable markers from the YTK (*4*). Co-expression of one or more “secretion booster” proteins can then be evaluated for enhanced secretion or surface display of a POI.

**Table 1.** Secretion booster proteins – endogenous yeast proteins that when overexpressed have previously been shown to enhance secretion of an heterologous protein of interest.

| Protein | Plasmid# | Step in Secretory Pathway | Short Description | Heterologous POI(s) Tested | References |
| --- | --- | --- | --- | --- | --- |
| Seb1p | pYSD509 | Translocation to ER | Sec61 translocon subunit | α-amylase | (59) |
| Sec65p | pYSD514 | Translocation to ER | SRP subunit | α-amylase | (60) |
| Srp14p | pYSD515 | Translocation to ER | SRP subunit | β-glucosidase, endoglucanase, $\alpha$ - amylase | (61) |
| Srp54p | pYSD519 | Translocation to ER | SRP subunit | β-glucosidase, endoglucanase, $\alpha$ -amylase | (61) |
| Ssa1p | pYSD518 | Translocation to ER | Cytosolic chaperone | β-glucosidase, Fab*, G-CSF* | (61-63) |
| Msn4p | pYSD507 | Translocation to ER | Stress response TF | VHH*, scFv* | (64) |
| Erj5p | pYSD504 | ER - protein folding | ER chaperone | VHH* | (64) |
| Sil1p | pYSD510 | ER - protein folding | ER chaperone | Fab* | (62) |
| Jem1p | pYSD526 | ER- protein folding | ER chaperone | rHA | (65) |
| Hac1p | pYSD506 | ER - protein folding | UPR transcription factor | α-amylase, rHA, VHH*, scFv* | (64-66) |
| Cwh41p | pYSD522 | ER - protein folding | Glucosidase | α-amylase | (67) |
| Mns1p | pYSD520 | ER - protein folding | Alpha-1,2-mannosidase | α-amylase | (60) |
| Ero1p | pYSD505 | ER- disulphide bond | Oxidoreductase | α-amylase; scTCR; scFv | (60, 68) |
| Pdi1p | pYSD508 | ER- disulphide bond | Protein disulfide isomerase | β-glucosidase, endoglucanase, $\alpha$ - amylase, G-CSF | (61, 63, 69) |
| Erv2p | pYSD513 | ER- disulphide bond | Sulfhydryl oxidase | α-amylase | (60) |
| Erv29p | pYSD503 | ER to Golgi | COPII cargo receptor | α-amylase | (70) |
| Sec16p | pYSD523 | ER to Golgi | COPII assembly scaffold | endoglucanase, $\alpha$ -amylase, glucan-1,4- α-glucosidase | (71) |
| Sed5p | pYSD525 | ER to Golgi | t-SNARE protein | β-glucosidase, cellobiohydrolase | (72) |
| Cog5p | pYSD517 | Golgi | Golgi COG complex | α-amylase | (70) |
| Sec4p | pYSD521 | Secretory Vesicles | Rab GTPase | β-glucosidase, α-amylase | (59, 73) |
| Sso1p | pYSD511 | Secretory Vesicles | t-SNARE protein | α-amylase, β-glucosidase, | (72-74) |
| Sso2p | pYSD512 | Secretory Vesicles | t-SNARE protein | α-amylase | (74) |
| Swa2p | pYSD516 | Vacuole Sorting | Clathrin uncoating factor | α-amylase | (60) |
| Cys4p | pYSD524 | Protein Synthesis | Cystathionine beta-synthase | α-amylase | (60) |

References and heterologous proteins mentioned are representative examples for a given secretion boosting protein and is not intended to be an exhaustive list. Asterisks indicates examples reported in *K. phaffii* rather than *S. cerevisiae*. Abbreviations: SRP = Signal recognition particle; VHH = variable heavy domain of heavy chain (camelid antibody); rHA = recombinant human albumin; G-CSF = Granulocyte colony stimulating factor; UPR = Unfolded Protein Response; COPII = coat protein complex II; SNARE = soluble NSF attachment protein receptor; scFv = single-chain variable fragment of antibody; COG = Conserved oligomeric Golgi complex; scTCR = single chain T-cell receptor

## CONCLUSIONS

With the additions described here the YSD toolkit now has eight different surface display anchors, most of which are available with three different epitope tags. Two options are available for C-terminal fusion of the POI to the anchor and six for N-terminal fusion. This represents a comprehensive set of surface display parts. In addition, sixteen different SPs (*25*) can be combined with the six N-terminal fusion anchors. Thus, in total, the toolkit allows 98 different coding sequences to be generated for a given POI in order to optimize its surface display. While robotic platforms would allow this number of individual expression plasmids to be generated, another option would be to combine the anchor and SP parts during the assembly of expression cassettes, thereby generating a small library of expression plasmids. This library could be screened by FACS for example to select the construct(s) that best display the POI. Similarly, for a secreted POI, expression cassettes with up to sixteen different SPs could be combined with expression cassettes for the twenty-four secretion booster proteins presented here. This could be done either by co-transformation of single transcriptional unit POI and secretion booster plasmids into yeast, or by generating a combinatorial library of multigene expression plasmids through golden gate assembly. The fluorescence complementation functionality that we have added to the toolkit should facilitate high throughput screening of such combinatorial libraries. Of course, the panel of secretion booster proteins can also be tested for enhancement of cell surface display of a POI. Overall the expanded YSD2.0 toolkit enables combinatorial screening of heterologous POI and secretion boosting expression constructs, potentially facilitating *Design of Experiments* (DoE) strategies for efficient, systematic exploration of design space (*75*).

The examples of surface display applications presented here showcase the utility of the YSD toolkit for co-display of up to three different proteins and to display affinity reagents that can be used to identify and characterize protein:protein interactions. The co-display aspect should be applicable to the generation of other whole cell biocatalysts besides the PET degradation example provided here. In addition, co-display of multiple (non-enzymatic) proteins may have applications in other bioengineering contexts involving yeast. The use of α-GFP Nb displaying yeast as a highly cost-effective tool to interrogate protein interactions has the potential to be broadly adopted. This concept can likely be expanded to the surface display of antibodies and other affinity reagents the sequences for which are increasingly becoming available (*76–78*). If successful surface display of such proteins can be achieved, then one has an essentially limitless source of the affinity reagent that can be expanded on demand to the scale required for a given experiment and shared with collaborators and the broader scientific community.

## METHODS

### Chemicals, molecular biology reagents

All chemicals were obtained from Merck unless stated otherwise. The MoClo-YTK plasmid kit was a gift from John Dueber (Addgene kit # 1000000061). A plasmid containing GFP1-10 (pHHYTK113) was a gift from Johannes Herrmann (Addgene plasmid # 200144). Enzymes used were from New England Biolabs and Fisher Scientific. Single stranded oligonucleotides and double stranded DNAs (gBlocks) were ordered from Integrated DNA Technologies.

### Strains and growth conditions

The *S. cerevisiae* strain used was BY4741 (MATa his3Δ1 leu2Δ0 met15Δ0 ura3Δ0). For selection of transformed plasmids, cells were grown at 30 °C in minimal synthetic media or agar plates containing 6.8 g/L yeast nitrogen base (Merck #Y0626) without amino acids, 5 g/L dextrose, and a complete supplement mixture of amino acids minus leucine or minus leucine and histidine (MP Biomedicals). Yeast transformation was performed using a standard protocol as previously described (*25*). *E. coli* DH5α cells were used for all cloning steps and were grown at 37 °C on Lysogeny Broth (LB) agar plates or in liquid LB media with shaking at 200 rpm. 25 μg/mL chloramphenicol, 30 μg/mL kanamycin or 50 μg/mL carbenicillin were added as appropriate.

### MoClo DNA assembly reactions and part generation

Protein encoding parts were designed to be compatible with the MoClo Yeast Toolkit (YTK) (*4*), with the exception of the overhang between the custom 3a’ and 3b’ parts (*25*). Smaller parts were generated by annealing and extending complementary oligonucleotides using Klenow polymerase. Larger parts were generated by PCR amplification from yeast genomic DNA, plasmids or synthetic DNA fragments. Golden gate assembly reactions and verification of constructs was performed as previously described (*25*). To generate anchors for amino terminal fusion of POIs, sequences coding for a flexible linker (GGGGSGGGGS), an epitope tag, a TEV protease cleavage site and the anchor protein were created as MoClo YTK type 4a parts (according to the nomenclature of Lee et al. (*4*)). The N-terminally deleted Aga1 anchor (Aga1ΔN181) comprised amino acids 182-544 of Aga1p (YNR044W) (*26*). The Aga2 anchors for carboxyl terminal fusion of POIs were created as a MoClo YSD type 3a’ parts (as defined in (*25*)) encoding an epitope tag after the Aga2p signal peptide (after amino acid 18) and a flexible linker sequence (GGGGSGGGGS) at the C-terminus of Aga2p (YGL032C). The Pir2 anchor was created as a MoClo YSD type 3a’ part with a flexible linker sequence (GGGGSGGGGS), an epitope tag and a TEV protease cleavage site located at the C-terminus of full length Pir2p (YJL159W). Coding sequences for LCC-WT (*36*), LCC-ICCG (*37*), HFB1 (*79*), MHETase (*39*), anti-HA scFV (*48*), Protein A ZZ-domain (*80*) and PDZ-binding motifs (*28, 46*) were created as MoClo YSD type 3a’ parts. GFP11 and GFP11-His^6^ were encoded as type 4a parts with a flexible linkers (SGSGGGS and GGSG) preceding the GFP11 and His^6^ sequences respectively. pET23b-15F11-HA-mEGFP-6xHis was a gift from Tim Stasevich (Addgene plasmid # 129593). Note that the α-GFP Nb construct used in this study contains an C-terminal HA epitope in addition to any tag present in the anchor protein. A part encoding the anti-GFP Nb without this HA tag is also provided (pYSD069). Yeast expression plasmids with CEN6/ARS4 origins of replication and *LEU2, URA3* or *HIS3* selectable markers were assembled using parts from the YTK and YSD toolkits as summarised in Supporting Information File S1. Double stranded oligonucleotides for the liprin-α1 and erbin PDZ binding motifs and PCR products for LCC-WT and -ICCG were assembled directly into expression plasmids without first creating entry vector constructs. For surface display of Aga2p, Aga1p was co-expressed from the plasmid pYSD300.

### Bacterial protein expression and purification

The following vectors were used for protein expression: pYTK001 (*4*) for GFP, pET24d (Merck/Novagen) for His^6^-GFP1-10, pRSF-Duet (Merck/Novagen) for Erbin-PDZ-citrineYFP-His^6^ and citrineYFP-His^6^, pBAD (ThermoFisher Scientific) for HA-tagged citrineYFP-His^6^ and FLAG-tagged citrineYFP-His^6^. *E. coli* DH5α was used for constitutive GFP expression, while *E. coli* BL21 (Merck/Novagen) was used for all other proteins.

Protein expression was induced for 4 hours in mid log phase cultures using 0.1 % arabinose (pBAD vector) or 0.1 mM isopropyl-β-D-thiogalactopyranoside (pET24d and pRSF-Duet vectors). Bacterial cell pellets were resuspended in bacterial lysis buffer (PBS, 0.2 % Triton X100, 20 mM β-mercaptoethanol, 1 mM phenylmethylsulfonyl fluoride (PMSF)) supplemented with 0.1 mg / mL lysozyme, and 0.1 mg/mL DNAse. Approximately 1 mL lysis buffer was used per 20 mL of culture volume. Following a 30 minute incubation on ice, cells were sonicated and the lysed cells were centrifuged at 16,100 x g for 30 minutes at 4 °C to obtain the soluble cell lysate.

In some cases, this lysate was used directly for flow cytometry and immunoprecipitation experiments. In other cases, to obtain purified proteins, cell lysates were incubated with Ni-NTA agarose beads (Neo Biotech), washed in 20 mM Tris pH 8, 500 mM NaCl, 20 mM β-mercaptoethanol, 5 mM imidazole, 0.1 % Triton X100 and eluted in 200 mM imidazole pH 7. Purified Erbin-PDZ-citrineYFP-His^6^ and citrineYFP-His^6^ were dialysed into a buffer containing 100 mM Tris pH 7.5, 50 mM NaCl, 1 mM dithiothreitol (DTT). For GFP1-10, which was largely insoluble, the pellet obtained after cell lysis was solubilized in 8 M urea, 0.5 M NaCl, 50 mM KH_2_PO_4_/K_2_HPO_4_ pH 8, 5 mM imidazole, 0.1 % Triton, purified using Ni-NTA agarose and eluted in 6 M urea, 200 mM imidazole pH 7. Purified GFP1-10 was dialysed into TNG buffer (100 mM Tris-HCl (pH 7.5), 150 mM NaCl, 10% (vol/vol) glycerol) and supplemented with 5 mM DTT prior to use in fluorescence complementation assays.

### Mammalian protein expression

HEK (Human Embryonic Kidney) 293 cells (ATTC), cultured under standard conditions, were transfected with mammalian expression constructs using calcium phosphate precipitation. GFP-LNX1 and FLAG-Liprin-α1 constructs were described previously, while FLAG-NUMB and FLAG-ZNF24 were prepared by cloning the coding sequences for mouse NUMB and human ZNF24 in the pCMV-N-FLAG vector (*46*). Transfected HEK293 cells were washed once with PBS, resuspended in mammalian cell lysis buffer (10 mM Tris/Cl pH 7.5, 150 mM NaCl, 0.5 mM EDTA, 0.5% NP40 detergent, 1 mM PMSF) and incubating on ice with occasional pipetting for 20 minutes. 75 μL lysis buffer was used per well of a six well culture dish and 200 uL per 10 cm dish. The lysed cells were centrifuged at 16,100 x g for 30 minutes at 4 °C and the supernatant diluted with three volumes of wash buffer (10 mM Tris/Cl pH 7.5, 150 mM NaCl, 0.5 mM EDTA).

### Flow cytometry

For flow cytometry applications, 2 mL yeast cultures were grown in 13 mL tubes, in appropriate selective synthetic media at 30 °C overnight, shaking at 200 rpm. Yeast cell density was measured. Approximately 1×10^6^ cells (assuming 1 OD_600_ ≈ 1.5 × 10^7^ cells/mL) were pipetted into each microcentrifuge tube and centrifuged for 3 minutes at 3500 × g. The supernatant was removed and cells were resuspended in 100 μL of flow cytometry buffer (20 mM HEPES pH 7.5, 150 mM NaCl, 0.1% (w/v) bovine serum albumin, 5 mM maltose). 0.5 μg of anti-epitope tag antibody and/or 20-30 μL of bacterial cell lysate or 50-100 μL of mammalian cell lysate was then added and cells incubated for 15 minutes at room temperature when using only antibodies or 4 °C when using cell lysates. After incubation, yeast cells were pelleted again, resuspended in 500 μL of flow cytometry buffer and analysed by flow cytometry on an Accuri C6 instrument (BD Biosciences). An exception to this was when flow cytometry was used to detect interactions of GFP-tagged LNX1. In this case after incubation with cell lysates the yeast cells were pelleted, washed once with 500 μL flow cytometry buffer, incubated with anti-FLAG antibody as indicated above, pelleted again and resuspended in 500 μL flow cytometry buffer for analysis. The antibodies used for flow cytometry, obtained from Biolegend, were Alexa Fluor 488 mouse IgG1a anti-HA-tag (#901509), PE-Cyanine7 rat IgG2a anti-FLAG(DYKDDDDK)-tag (#637323) and Alexa Fluor 647 mouse IgG1a anti-c-Myc-tag (#626810). These were analysed using the FL1 (488 nm laser; 533/30 nm emission filter), FL3 (488 nm laser; 670 nm long pass emission filter) and FL4 (640 nm laser; 675/25 nm emission filter) detectors respectively. GFP and YFP were also analysed using the FL1 detector. An Alexa Fluor 488 conjugated donkey anti-rabbit (Jackson ImmunoResearch) and the above anti-c-Myc antibody were used to evaluate the capture of immunoglobulins by yeast displaying the Protein A ZZ domain. Unstained samples and stained samples of a strain not displaying the protein or tag of interest were used as controls. Analysis of flow cytometry data was performed using Flowreada (https://floreada.io) or FlowJo (https://www.flowjo.com) software.

### PET degradation assays and HPLC analysis

Amorphous PET film (6.3% crystallinity, 0.25□mm thick, Goodfellow Cambridge, #457-814-65, #ES30-FM-000145) was punched into 6 mm diameter discs with an average mass of 12 mg using a standard office paper punch. The film was washed in 0.5% Triton X-100 at 50□°C for 30□min, 10□mM Na_2_CO_3_ at 50□°C for 30□min, deionised water at 50□°C for 30□min and 70% ethanol for 5 min before air-drying. The film was then placed into a tube with 1□mL of buffer (either 100 mM glycine-NaOH pH 9 or 100 mM sodium phosphate buffer pH 8) containing 1 × 10^8^ of yeast cells for 72 h at 30 °C, 40 °C or 50 °C. The reaction was stopped by adding an equal volume of methanol, heating at 85□°C for 10□min, and separating the cells from the reaction with a 0.22 μm filter. 20 µL of each reaction was analysed by reversed-phase HPLC to measure TPA, MHET, and BHET production. HPLC was performed using an Agilent 1200 Series system (Agilent Technologies) equipped with a Zorbax SB-C8 column (4.6□×□150 mm, 5 µm). The column temperature was maintained at 22 °C. The mobile phase was solvent A (1% acetic acid in H_2_O) and solvent B (1% acetic acid in acetonitrile), at a flow rate of 1 mL/min. The analytes were eluted using a gradient over 25 min under the following conditions: solvent B content was increased from 5% to 44% over 13□min, then to 70% over 5 min, at which point the ratio was kept constant for 5□min, followed by decreasing solvent B to 5% over 1 min and keeping the ratio constant for 1 min. Detection wavelength was 240 nm (signal wavelength□=□240 nm with 4 nm bandwidth; reference wavelength□=□450 nm with 80 nm bandwidth). TPA (#100-21-0), MHET (#1137-99-1), and BHET (#959-26-2), all from Merck/Sigma-Aldrich, were dissolved in DMSO to generate standard curves of the relationship between area under the peak and the concentration of the standard. The standard curves were used to quantify detected TPA, MHET and BHET. The retention times were ∼7.2 min, ∼8.2 min and ∼8.7 min for TPA, MHET and BHET, respectively. All experiments were carried out in triplicate.

### GFP fluorescence complementation assays

10 mL cultures of yeast cells were grown in selective synthetic media supplemented with 0.85 % MOPS free acid, and 0.1 M dipotassium phosphate (both adjusted to pH 7). Once cultures reached an OD_600nm_ of between 2 and 2.4 they were centrifuged for 10 min at 4800 g. Secreted His-tagged proteins were captured from the remaining supernatant by adding 40 uL Ni-NTA agarose beads (Neo Biotech), incubating for 10 min at room temperature, centrifuging for 5 minutes at 1000 g and eluting in 120 uL of 200 mM imidazole (pH 7). A Greiner Bio-One black 96 well microplate (#655209) was prepared by adding 250 uL of blocking buffer (TNG, 0.5% bovine serum albumin (BSA)) to each well, incubating at room temperature for 10 min and then removing the blocking buffer. 60 uL of the eluted protein was combined with 100 uL of purified GFP1-10 and 40 uL of TNG buffer supplemented with 0.5% BSA in each well. mRuby2 and GFP fluorescence were measured immediately on a Tecan Infinite M200 plate reader using 540 nm (25 nm) / 575 nm (50 nm) and 465 nm (35 nm) / 520 nm (30 nm) excitation / emission filters respectively. The plate was incubated at room temperature and GFP fluorescence measured at 1.5, 6 and 24 hours. The plate was sealed between measurements to prevent evaporation. To account for small differences in cell density fluorescence measurements were normalized based on OD_600nm_ measurements of the whole cultures.

### Western blotting

Following SDS-PAGE proteins were transferred onto Immobilon-FL PVDF membranes (Thermo Fisher Scientific). Western blotting was performed using either rabbit anti-GFP (Abcam #AB290), mouse anti-FLAG (Merck/Sigma Aldrich #F3165) mouse anti-His tag (Genscript #A00186) or rabbit anti-NUMB (Abcam #14140) primary antibodies and IR700 or IR800 conjugated secondary antibody. Images were acquired on an Odyssey imaging system and analysed using Image Studio software (Li-Cor Biosciences).

### Immunoprecipitation using yeast

The plasmids used for experiments involving immunoprecipitation of GFP or GFP-tagged proteins were pYSD310 and pYSD315, which express either the α-GFP Nb or mRuby (as a negative control) fused to the HA-tagged 649 stalk anchor protein under control of the strong TEF1 promoter. Overnight cultures of yeast harbouring these plasmids were grown in selective synthetic media at 30 °C. Approximately 1 × 10^8^ cells were used per immunoprecipitation (assuming 1 OD_600_ ≈ 1.5 × 10^7^ cells/mL). Cells were centrifuged at 3500 x g for 5 min, resuspended in wash buffer (10 mM Tris/Cl pH 7.5, 150 mM NaCl, 0.5 mM EDTA) supplemented with 0.1% BSA, incubated for 5-10 min on ice and centrifuged again. Cells were then resuspended in either 25 uL of bacterial cell lysate plus 475 uL of wash buffer or 400-500 μL of mammalian cell lysate (prepared as described above). This was incubated for 30 minutes on a rotating mixer at 4 °C, then centrifuged at 4 °C and washed 3 times in wash buffer. Bound proteins were eluted by incubating the yeast cells for 5 min in one of the following: (1) 50 μL of 0.2 M glycine pH 2.5, (2) 60 μL of 0.2% SDS, 100 mM Tris pH 7.5, (3) 60 μL of 8 M Urea, (4) 60 μL of 2X SDS-PAGE sample buffer (100 mM Tris pH 6.8, 4% SDS, 20% glycerol, 10% β-mercaptoethanol, 0.01% bromophenol blue). Temperatures used for elution are indicated in the results section. Cells were pelleted by centrifugation and supernatants removed. The supernatant from low pH elutions were immediately neutralized by addition 10 μL of 1.5 M Tris pH 8.8. Supernatant samples were mixed with 3X SDS sample loading buffer, boiled and analysed by SDS-PAGE and western blotting.

To compare immunoprecipitation using yeast to magnetic beads a bacterial cell lysate from GFP expressing *E. coli* was prepared as described above except that mammalian cell lysis buffer was used and the cleared cell lysate was diluted with three volumes of wash buffer. This ensured that the buffer conditions matched those recommended for the ChromoTek GFP-Trap® magnetic agarose beads (Proteintech #gtma-20). Washing and elution was performed as described above for yeast cells, except that separation of GFP-Trap beads was performed using a magnetic tube rack rather than centrifugation.

### Plasmid availability

Full details of all MoClo plasmids used in this study are provided in Supporting Information File S1. Part plasmids in the pYTK001 entry vector with the coding sequences for anchor proteins, affinity reagents, GFP11 and secretion booster proteins, as well as a selection of expression constructs, will be submitted to Addgene and made available as version 2.0 of the Yeast Secretion/Display (YSD) plasmid kit (https://www.addgene.org/Paul_Young/). Fully annotated sequence files (in Genbank format) for the pYSD plasmids used in this study (that were not previously provided in O’Riordan et al. (2024) (*25*)) have been added as supporting information to this article (File S2). Sequences for plasmids that are included in the YSD 2.0 toolkit will also be available through the Addgene webpage for each plasmid (navigate to Resource Information -> Supplemental Documents).

## ABBREVIATIONS

α-GFP Nb: anti-GFP nanobody
AA: amino acid
BSA: bovine serum albumin
BHET: bis(2-hydroxy-ethyl) terephthalate
EG: ethylene glycol
ER: endoplasmic reticulum
FACS: fluorescence-activated cell sorting
FSC: forward light scattering
GFP: green fluorescent protein
GPI: glycosylphosphatidylinositol
GRAS: generally recognized as safe
HA: hemagglutinin
HFB1: hydrophobin 1
LCC: leaf-branch compost cutinase
MHET: monohydroxy□ethyl terephthalate
MoClo: modular cloning
PBS: Phosphate buffered saline
PET: poly(ethylene terephthalate)
POI: protein of interest
SD: standard deviation
SP: signal peptides
SRP: signal recognition particle
TPA: terephthalic acid
YSD: yeast secretion/display
YTK: yeast toolkit

## AUTHOR CONTRIBUTIONS

VJ, LE, and PY performed experiments and data analysis. NO’R developed the original YSD toolkit that forms the basis for this work. EM performed initial characterization of the LCC enzymes used in this study. PY and JH conceived and supervised the project.

## ACKNOWLEDGMENTS

We thank Rachel Breen and Dr. Anna Hogan in the School of Chemistry for their assistance with HPLC analysis. We are grateful to Joan Lenihan for providing transfected cell lysates. We thank Dr John Dueber, University of California, Berkeley for the MoClo yeast toolkit obtained through Addgene. We acknowledge the contributions of Jason Scully, Lia Teehan O’Connor, Ksenija Semjonova, Olivia Nix and Eimear O’Donoghue to the cloning of secretion booster genes. This work was supported by the Science Foundation Ireland AMBER Centre for Advanced Materials and BioEngineering Research (PY, EM, JH and VJ; Project Number 12/RC/2278_P2), the Irish Research Council through an Enterprise Partnership Postgraduate Scholarship to NO’R (Project ID: EPSPG/2020/307) and a Lilly Research Scholarship to LE.

## SUPPORTING INFORMATION

Supporting Figures and Tables that include additional data and information described in the text: (Figure S1) Characterization of new anchors added to the MoClo yeast surface display toolkit; (Figure S2) Optimization of conditions for immunoprecipitation using yeast displaying an a-GFP Nb; (Figure S3) Functionalization of yeast for analysis of protein:protein interactions using other affinity reagents.

A supporting file that provides: descriptions of part plasmids used and generated; descriptions of assembled expression plasmids; a list of pYSD plasmids that will be made available through Addgene.org (Supporting File 1) XLSX A supporting file with annotated sequences in GenBank format for the pYSD plasmids used in this study (that were not previously provided in O’Riordan et al. (2024) (*25*)). (Supporting File 2) ZIP

