## Supplementary Figures for "Yeast MoClo secretion and surface display toolkit 2.0: improvements and applications for analysis of protein-protein interactions and whole-cell biocatalysis"

**Figure S1**

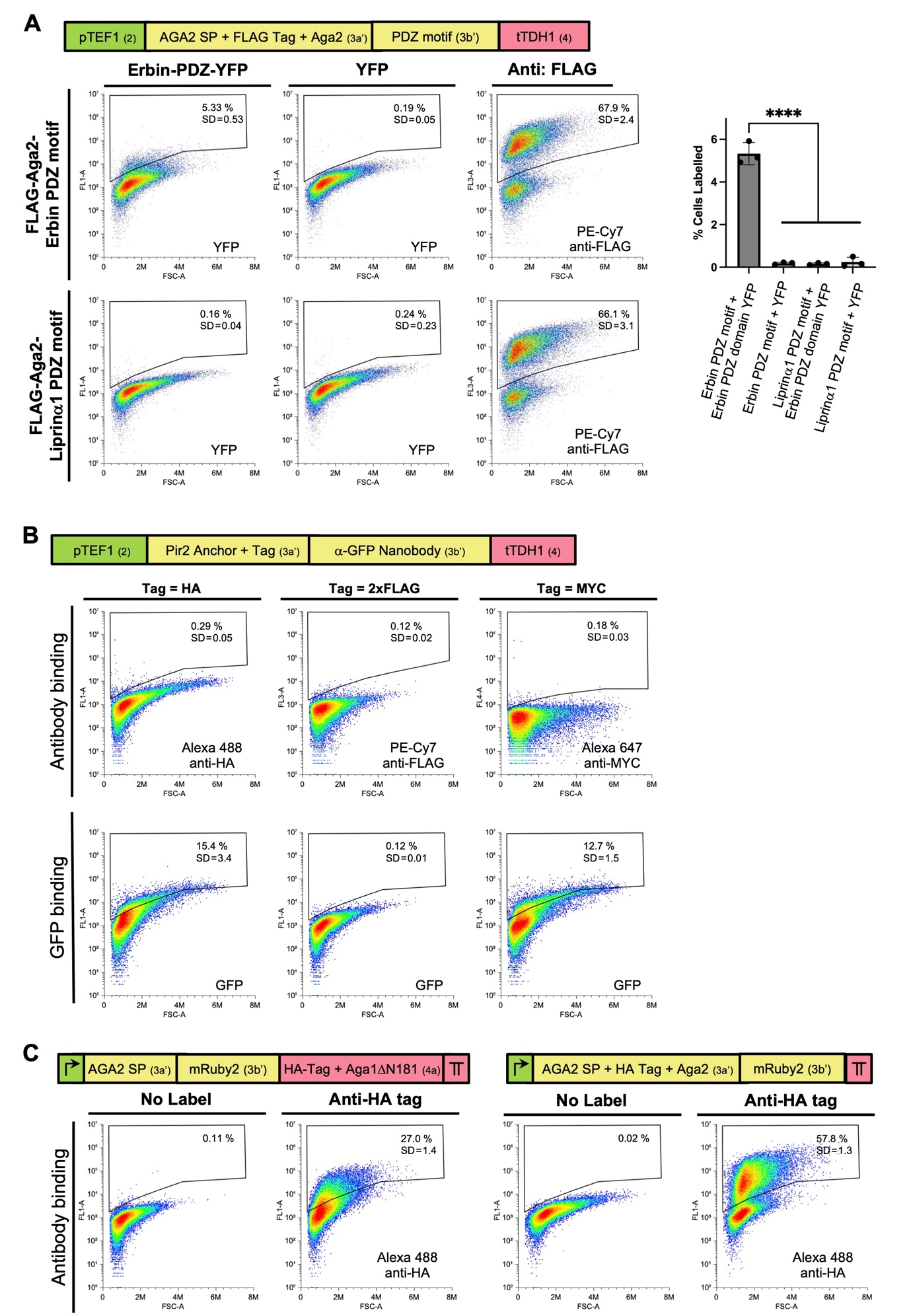

**Figure S1 Characterization of new anchors added to the MoClo yeast surface display toolkit**

**(A)** Detection of the interaction of a PDZ domain with a PDZ-binding motif displayed as a carboxyl terminal fusion to Aga2. Yeast strains were generated that displayed either a motif that is known to bind to the Erbin PDZ domain *(1)*, or an unrelated PDZ-binding motif from Liprin-α1 *(2)*, as carboxyl terminal fusions to FLAG-tagged Aga2. Yeast cells were incubated with either the Erbin-PDZ domain fused to YFP or YFP alone (as a negative control). Greater than 66% of cells are positively labelled using an anti-FLAG antibody, indicating efficient surface display of both PDZ-motifs. 5.33% of cells displaying the Erbin PDZ-binding motif are positively labelled with Erbin-PDZ-YFP versus 0.19% for YFP alone. Negligible binding of either Erbin-PDZ-YFP or YFP alone to cells displaying the Liprin-α1 PDZ-binding motif was observed. These results indicate very specific, though somewhat inefficient binding of the Erbin PDZ domain to a yeast surface-displayed carboxyl terminal PDZ motif. Quantification of binding from three independent experiments analysed by one-way ANOVA and Tukey’s multiple comparisons test is shown (right panel). Graph plots the mean % of YFP-labelled cells as individual data points with error bars representing SD. **** P < 0.0001 **(B)** Characterization of a Pir2 anchor for carboxyl terminal fusion of proteins of interest. Type 3a’ MoClo parts encoding Pir2 with each of three epitope tags were assembled into expression constructs that display the α-GFP Nb as a carboxyl terminal fusion. Surface display and functionality of the nanobody was assessed by flow cytometry using anti-epitope tag anti bodies as well as GFP binding. **(C)** Surface display of mRuby2 as an N-terminal fusion to Aga1 ΔN181 (left panel) and as a C-terminal fusion to Aga2 (right panel). Surface display was assessed by flow cytometry using an anti-HA tag antibody. Plots depict the relevant fluorescence signal (Y-axis) versus forward light scattering (X-axis). The mean percentage of positively stained cells from 3-4 independent experiments is shown on a representative plot with the standard deviation (SD) indicated. Schematics of expression constructs indicate the promoter, terminator and signal peptide (SP) sequences used as well as part types (in brackets). The promoter and terminator in (C) are the same as in (A) and (B) but are shown as symbols for brevity.

**Figure S2**

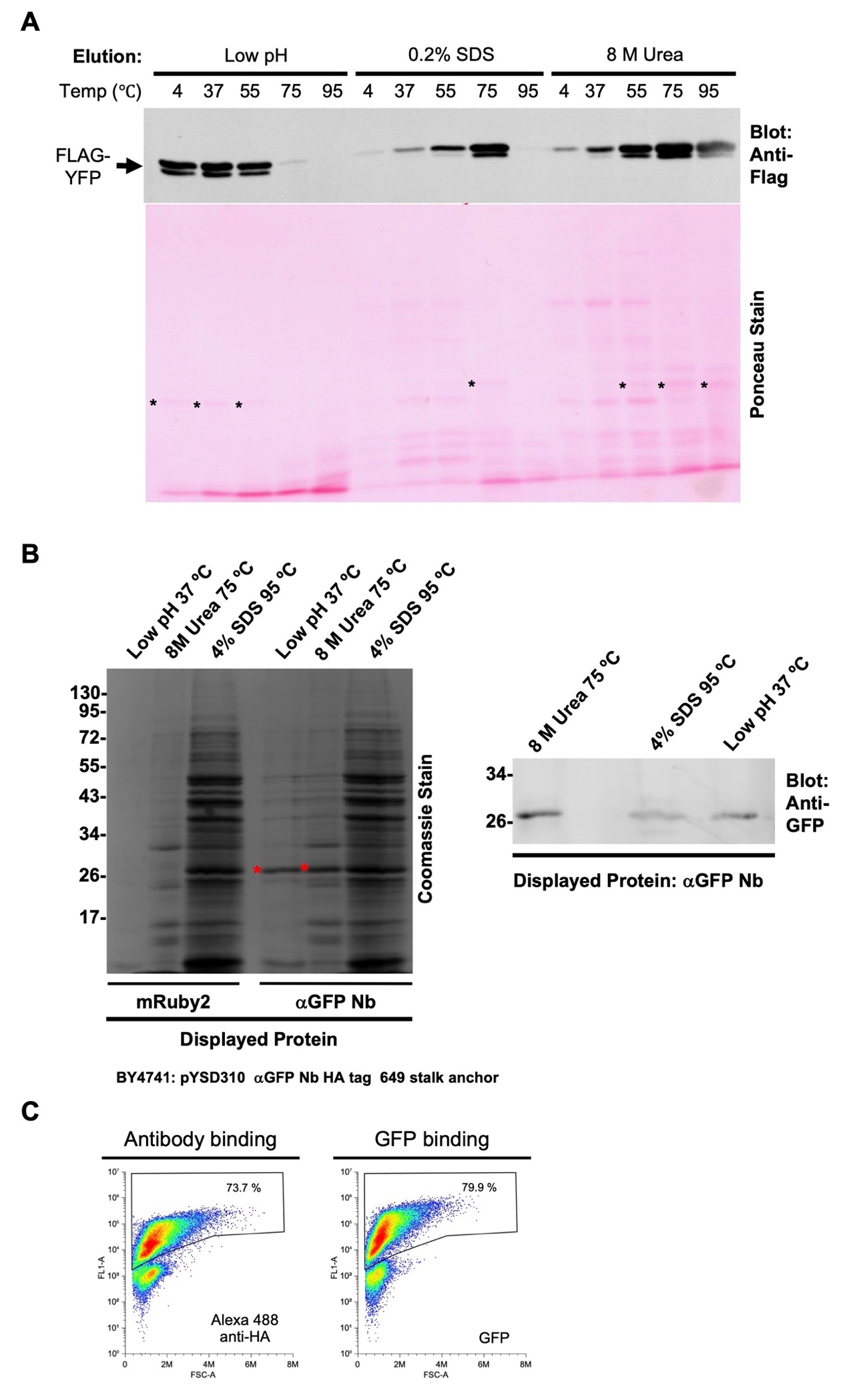

**Figure S2 Optimization of conditions for immunoprecipitation using yeast displaying an α-GFP Nanobody**

**(A)** Optimization of elution conditions. FLAG-tagged yellow fluorescent protein (YFP) was immunoprecipitated from an *E. coli* cell lysate using 1 x 10^8^ yeast cells displaying an α-GFP Nb (α-GFP-Nb yeast). After washing, bound proteins were eluted either by incubating for 5 minutes in either 0.2 M Glycine pH 2.5 (low pH), 0.2% SDS / 100 mM Tris pH 7.5 or 8 M Urea at the indicated temperatures. Eluates were analysed by Ponceau staining and western blotting to detect total protein and FLAG-YFP respectively. **(B)** Specific immunoprecipitation of GFP from a lysate of GFP-expressing *E. coli* cells using 1 x 10^8^ yeast cells displaying either an α-GFP Nb or mRuby2 (as a negative control). After washing, bound proteins were eluted either by incubating for 5 minutes in 0.2 M Glycine pH 2.5 at 37 ºC (low pH), 8 M Urea at 75 ºC or SDS-PAGE sample buffer (4% SDS) at 95 ºC. Eluates were analysed by Coomassie staining and western blotting to detect total protein and GFP respectively. GFP is visible by Coomassie staining in the low pH and 8M Urea elutions (red asterisks), while for the 4% SDS/95 ºC elution the GFP band is obscured by the release of large amount of yeast proteins. Western blotting shows elution with 8 M Urea at 75 ºC to yield slightly higher amounts of GFP than the other two elution conditions. **(C)** Surface display and binding activity of α-GFP Nb yeast following medium-term storage at 4 ºC. Flow cytometry analysis was performed on a liquid culture of yeast cells harbouring the pYSD310 plasmid that had been grown overnight in selective media and then stored for two weeks in a fridge.

**Figure S3**

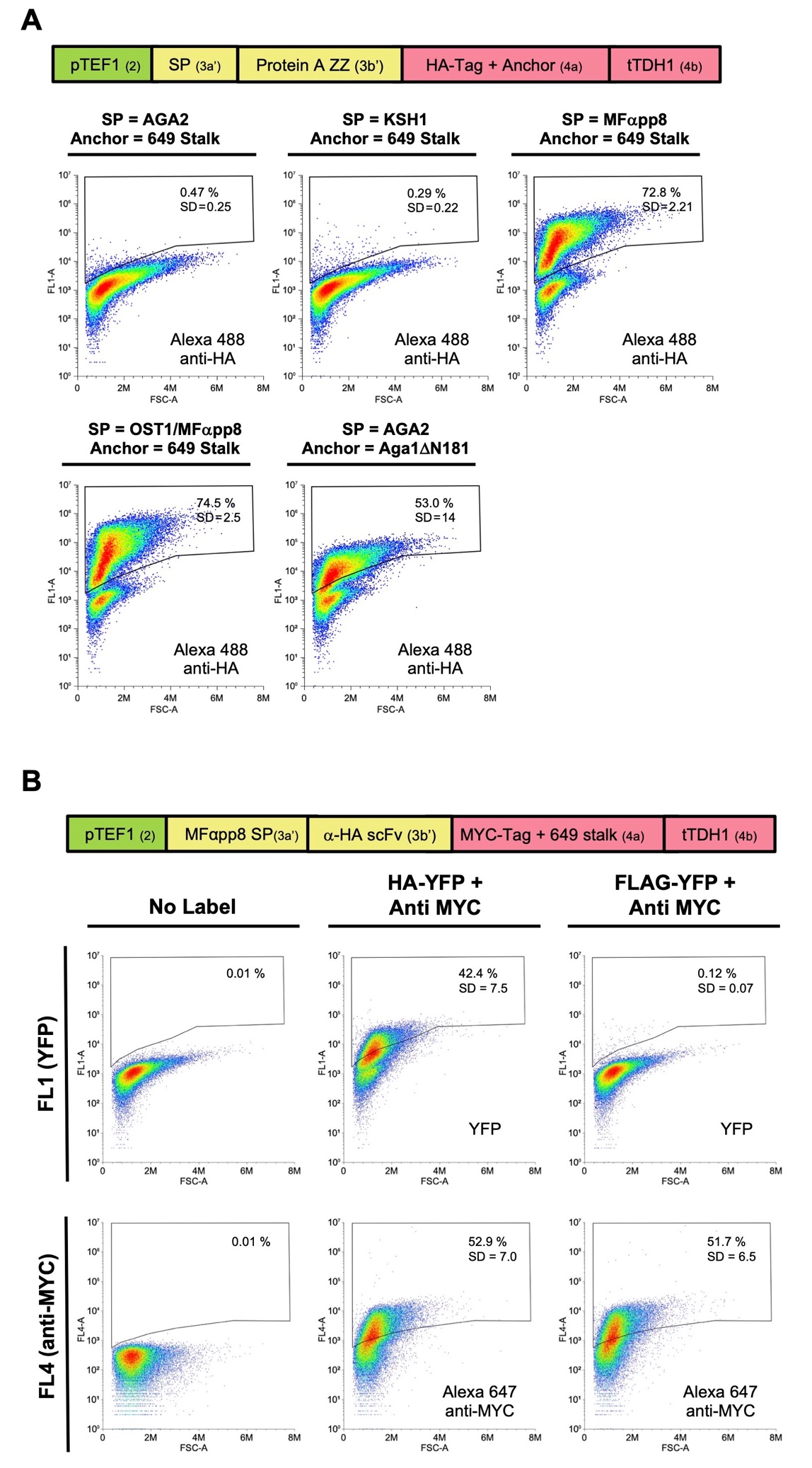

**Figure S3 Functionalization of yeast for analysis of protein:protein interactions using other affinity reagents.**

**(A)** Screening of SPs and anchor proteins to achieve efficient surface display of the Protein A derived ZZ immunoglobulin binding domain. Yeast cells transformed with expression constructs containing the indicated SP / anchor protein combinations were analysed by flow cytometry using an anti-HA antibody to evaluate the extent of ZZ domain surface display.

**(B)** Yeast surface display of an anti HA-tag scFv and its specific binding to HA-tagged proteins. Yeast cells displaying a MYC-tagged anti HA-tag *Frankenbody* scFv *(3)* were incubated with lysate from *E. coli* cells expressing either HA-tagged YFP or FLAG-tagged YFP (as a negative control). Surface display of the scFv was assessed by flow cytometry using an Alexa-647-conjugated anti-MYC antibody, while binding to the target protein was indicated by YFP fluorescence. Unlabelled cells are shown to verify the specificity of anti-MYC antibody and YFP binding. Greater than 50% of cells expressing the scFv are positively labelled using the anti MYC antibody, indicating efficient surface display. 42.4% of cells are positively labelled with HA-tagged YFP versus 0.12% for FLAG-tagged YFP, indicating specific binding of the scFv to HA-tagged proteins on the cell surface. The mean percentage of stained cells from four independent experiments is shown on a representative plot with the standard deviation (SD) indicated. Schematics of the MoClo-generated expression constructs are shown indicating the promoter, terminator, SP and displayed protein as well as part types (in brackets).
